# PARP activity is essential for retinal photoreceptor survival in the human homologous *Rho*^I255del^ mouse model for autosomal dominant retinitis pigmentosa

**DOI:** 10.1101/2025.08.21.671267

**Authors:** Yu Zhu, Azdah Hamed A Fallatah, Kangwei Jiao, Mathias W. Seeliger, François Paquet-Durand

## Abstract

Retinitis Pigmentosa (RP) is a group of rare, inherited, neurodegenerative diseases of the retina that primarily affect rod photoreceptors. The initial loss of rods is followed by a secondary cone photoreceptor degeneration and eventually legal blindness. Despite several attempts, RP still remains essentially untreatable. In recent years, inhibition of poly(ADP-ribose)polymerase (PARP) has been proposed as a potential therapeutic strategy of autosomal-recessive RP, based on promising work in preclinical animal models. However, the effects of PARP inhibitors in autosomal-dominant RP are still largely unknown.

Here, we employed a novel, human-homologous rhodopsin-mutant *Rho*^I255del/+^ mouse model for autosomal dominant RP to assess the impact of different PARP inhibitors on the progression of photoreceptor degeneration. The PARP inhibitors used –olaparib, saruparib, INO1001, and nicotinamide– target different PARP isoforms, and their potentially differential effects were evaluated in organotypic retinal explants cultivated under entirely defined conditions. Readouts comprised *in situ* activity assays for PARP and calpain-type proteases, the TUNEL assay for cell death, as well as immunostaining for activated calpain-2, activated caspase-3, rhodopsin, and cone arrestin-3.

Unexpectedly, and in contrast to previous findings in animal models for recessive RP, all of the included PARP inhibitors led to a marked and dose-dependent rod photoreceptor toxicity in the *Rho*^I255del^ model for dominant RP. Furthermore, this effect appeared to be independent of rhodopsin expression. On the other hand, both cone photoreceptors and inner retinal neurons were apparently unaffected by PARP inhibition.

The present study thus demonstrates the importance of PARP activity for rod photoreceptor viability in a dominant rhodopsin mutant, highlights the need for a deeper understanding of the mechanisms underlying photoreceptor degeneration in different RP forms, and cautions against the indiscriminate use of PARP inhibitors for the treatment of RP.

## Introduction

Retinitis pigmentosa (RP) refers to a group of inherited retinal degenerations (IRD), in which mutations in a large variety of genes cause primary rod photoreceptor death. Since rods are responsible for dim light vision, this initially causes night blindness. Subsequently, cone photoreceptors also succumb to degeneration, eventually resulting in complete blindness (Kaplan et al., 2017). While about 50-60% of RP cases show autosomal recessive inheritance, 30-40% of patients suffer from autosomal dominant disease (Bunker et al., 1984; Grøndahl, 1987; Novak-Lauš et al., 2002). Within the autosomal dominant group, approx. 25 % of cases are caused by mutations in the rhodopsin (*RHO*) gene (Hartong et al., 2006). To this day, no effective treatments exist for most forms of RP.

Numerous animal models are available to study RP treatment development (Arango-Gonzalez et al., 2014; Power et al., 2020). Across many of these disease models, both for recessive and for dominant disease, an over-activation of poly(ADP-ribose) polymerase (PARP) has been found (Arango-Gonzalez et al., 2014; Jiao et al., 2016; Kaur et al., 2011). The 17-member PARP family of enzymes play an important role in promoting DNA repair and regulating gene transcription. Notably, the synthesis of poly(ADP-ribose) (PAR) polymers is seen as a crucial factor that helps unwinding chromatin structure and facilitating DNA access of repair enzymes and transcriptional machinery (Bai, 2015). Paradoxically, excessive activation of PARP is linked to cell death (Park et al., 2022), including in retinal photoreceptors (Arango-Gonzalez et al., 2014), a phenomenon that may be related to an excessive consumption of nicotinamide adenine dinucleotide (NAD^+^). Accordingly, PARP inhibitors, such as olaparib (brand name Lynparza) or INO1001, have previously been reported to provide photoreceptor neuroprotection in different models for recessive RP *in vitro* (Dong et al., 2023; Sahaboglu et al., 2016; Yan et al., 2022) and *in vivo* (Sahaboglu et al., 2016). However, the efficacy of PARP inhibitors in protecting photoreceptors in dominant forms of RP is still largely unclear.

Here, we employed the recently generated human homologous *Rho*^I255del^ knock-in mouse model for dominant RP (Bowen Cao et al., 2024; Zhu, Kumar, et al., 2025; Zhu, Peiroten, et al., 2025), to test different PARP inhibitors for potential neuroprotective effects. Surprisingly, even though also in *Rho*^I255del^ photoreceptors high PARP activity was shown to be associated with photoreceptor degeneration (Bowen Cao et al., 2024), we found PARP inhibitors to exert strong negative effects on *Rho*^I255del^ rod photoreceptors. These results point to possible differences in degenerative mechanisms between recessive and dominant RP and also suggest that PARP inhibition as a potential therapeutic strategy may need to be tested and tailored to individual disease genotypes.

## Materials & Methods

### Animals

*Rho*^l255del/+^ (*Rho*^l255d/+^) mice carry a human homologous gene defect that results in the deletion of one of the two isoleucine residues located in positions 255 and 256 of the rhodopsin protein (Bowen Cao et al., 2024). The study employed heterozygous C57BL/6J *Rho*^l255del/+^ mice that were generated by crossing homozygous *Rho*^l255del/I255del^ with C57BL/6J *Rho*^+/+^ wild-type (WT) animals. Animals were housed in a specified pathogen free (SPF) facility, under standard white cyclic lighting, had free access to food and water, and were used irrespective of gender. All efforts were made to minimize the number of animals used. To limit the pain and suffering of animals, and to reduce the overall numbers of animals required, all experiments were conducted using *in vitro* organotypic retinal explant cultures. A total of 88 animals (166 retinal explants) were used for the study.

### Organotypic retinal explant cultures

Retinal explantation and cultivation under sterile conditions followed previously published protocols (Belhadj et al., 2020). In brief, *Rho*^l255del/+^ and WT animals were sacrificed at postnatal day (P) 12 by use of CO_2_ asphyxiation followed by cervical dislocation. The eyes were enucleated and incubated in basal R16 retinal culture medium (BM; 07491252A, Gibco, Paisley, UK) for 5 minutes (min) at room temperature (RT). To facilitate explantation of the neuroretina with attached retinal pigment epithelium (RPE), the sclera was predigested for 17 min with 0.12% proteinase K (39450, MP Biomedicals, Irvine, CA, USA) in R16 BM. To inactivate proteinase K, the eyes were incubated in R16 BM containing 20% fetal bovine serum (FBS; F7524, Sigma) for 5 min at RT. Under a stereoscope, the eyes were then dissected in fresh R16 BM, the anterior segments, sclera, choroid were removed, and the optic nerve was cut, leaving only the neuroretina with its RPE attached. To flatten the retina, four incisions were made, resulting in a four-leaf clover shape. The retinal explant was then transferred to a 6-well culture plate with polycarbonate membrane inserts (Corning Life Sciences, Corning, NY, USA), with the RPE facing the membrane. Retinal explant cultures were maintained in 1.2 mL of complete R16 medium (CM) with supplements, free of serum and antibiotics (Belhadj et al., 2020), in a sterile, humidified incubator at 37 °C with 5% CO_2_.

After 48h (*i*.*e*. at an age equivalent to P14), retinal explants were separated into non-treated (NT) control, exposed only to CM and dimethyl sulfoxide (DMSO; 0.01, 0.1, and 1%), and treatment groups exposed to different PARP inhibitors dissolved in DMSO and CM. The PARP inhibitors olaparib, saruparib, and INO1001 were used at 0.1, 1, and 10 µM concentrations, while nicotinamide (NAM) was used at 20, 200, 1000, and 2000 µM. For chemical structures and specificities of the PARP inhibitors used here see Table 1. The culture medium was changed every second day, and culturing ended at either P18 or P20 (Fig.1).

**Table 1.**
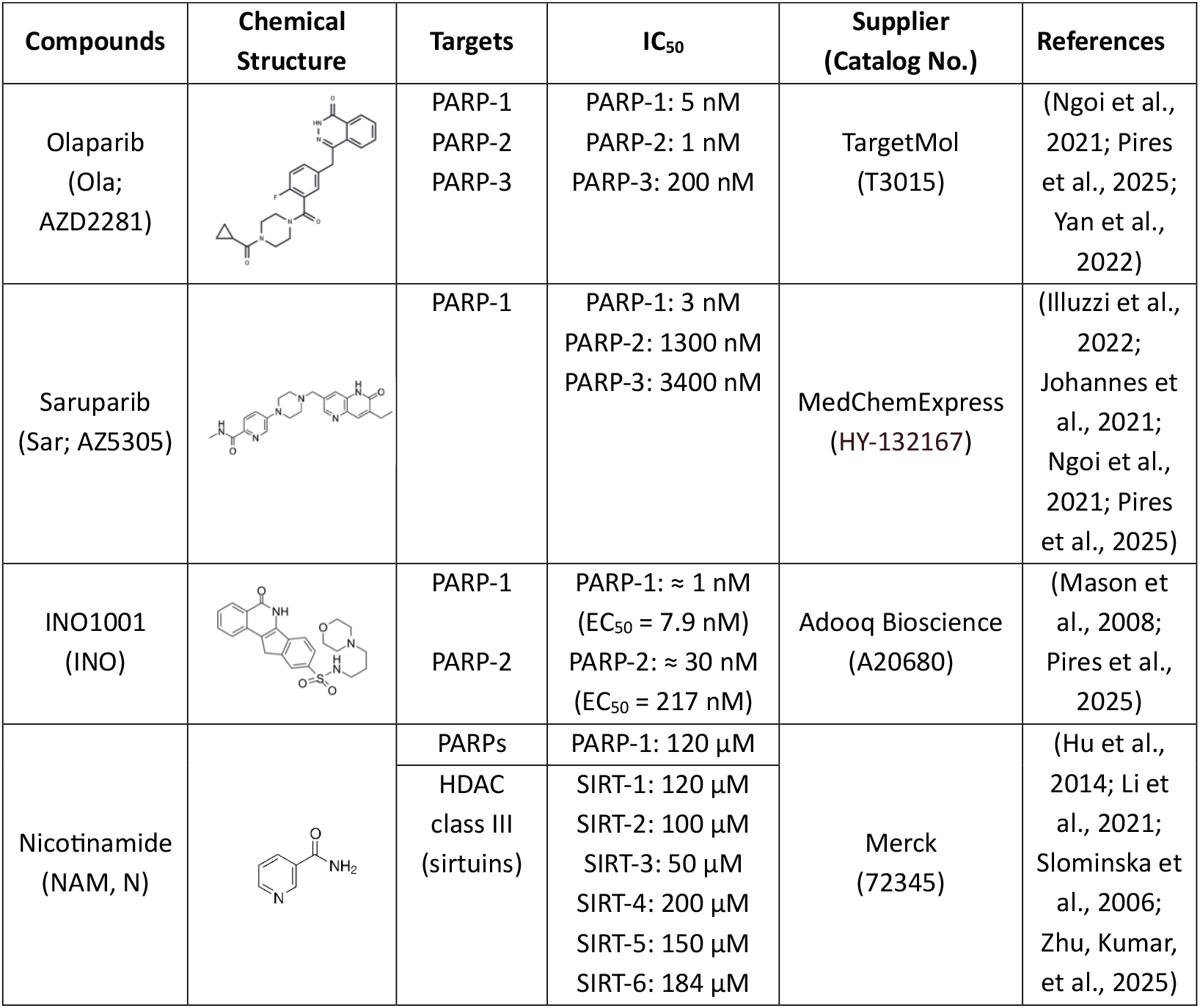
Overview of different PARP inhibitors used in this project. Information on different PARP inhibitors, including chemical structures, targets, IC_50_-values, providers, and corresponding references.

| Compounds | Chemical Structure | Targets | IC<sub>50</sub> | Supplier (Catalog No.) | References |
| --- | --- | --- | --- | --- | --- |
| Olaparib (Ola; AZD2281) |  | PARP-1<br>PARP-2<br>PARP-3 | PARP-1: 5 nM<br>PARP-2: 1 nM<br>PARP-3: 200 nM | TargetMol (T3015) | (Ngoi et al., 2021; Pires et al., 2025; Yan et al., 2022) |
| Saruparib (Sar; AZ5305) |  | PARP-1 | PARP-1: 3 nM<br>PARP-2: 1300 nM<br>PARP-3: 3400 nM | MedChemExpress (HY-132167) | (Illuzzi et al., 2022; Johannes et al., 2021; Ngoi et al., 2021; Pires et al., 2025) |
| INO1001 (INO) |  | PARP-1<br>PARP-2 | PARP-1: ≈ 1 nM (EC<sub>50</sub> = 7.9 nM)<br>PARP-2: ≈ 30 nM (EC<sub>50</sub> = 217 nM) | Adooq Bioscience (A20680) | (Mason et al., 2008; Pires et al., 2025) |
| Nicotinamide (NAM, N) |  | PARPs<br>HDAC class III (sirtuins) | PARP-1: 120 μM<br>SIRT-1: 120 μM<br>SIRT-2: 100 μM<br>SIRT-3: 50 μM<br>SIRT-4: 200 μM<br>SIRT-5: 150 μM<br>SIRT-6: 184 μM | Merck (72345) | (Hu et al., 2014; Li et al., 2021; Slominska et al., 2006; Zhu, Kumar, et al., 2025) |

**Figure 1:**
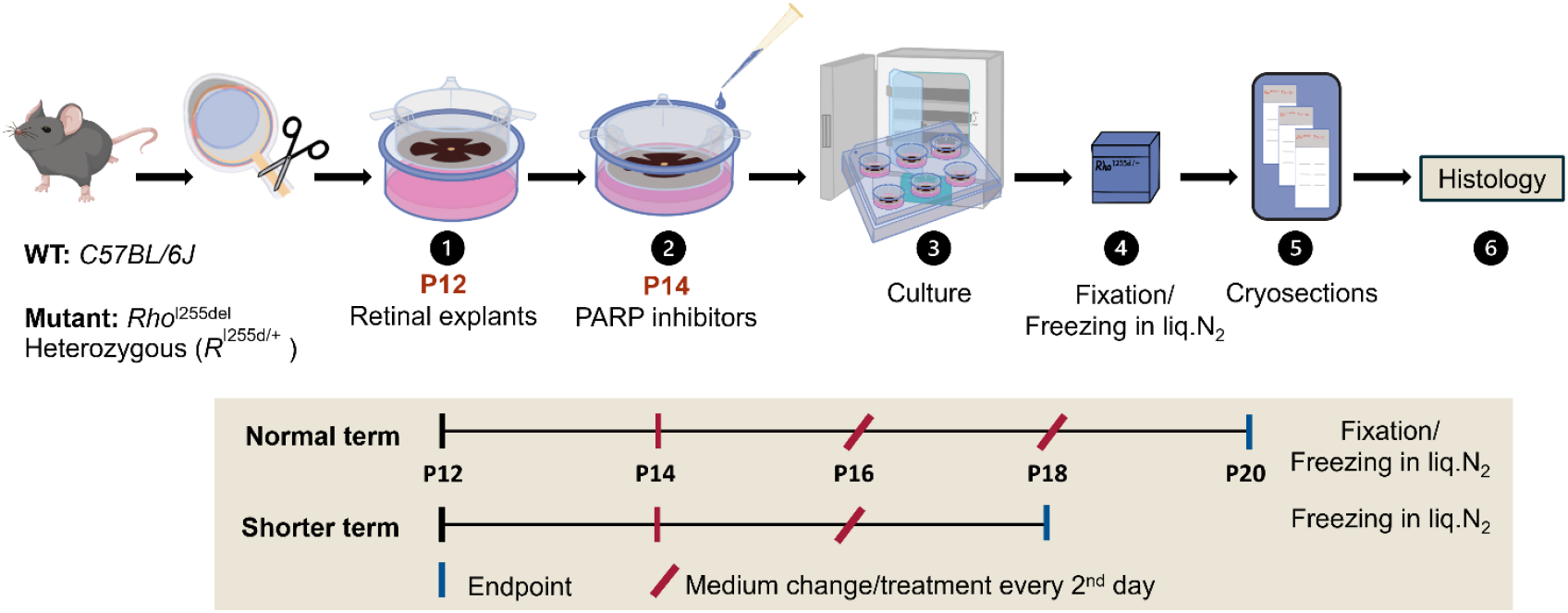
Schematic overview of experimental paradigms. Organotypic retinal explant cultures were derived from either wild-type (WT) or *Rho*^I255d/+^ animals at post-natal day (P) 12 and cultured until P18 or P20. The culture medium was changed every second day, and treatment with PARP inhibitors olaparib, saruparib, INO1001, and nicotinamide (or dimethyl sulfoxide (DMSO) control) was applied from P14 onwards. Cultures ended at P18 or P20, either by 4% paraformaldehyde (PFA) fixation (fixed tissue) or by immediate freezing in liquid N_2_(unfixed tissue).

### Histology

Retinal explants were fixed with 4% paraformaldehyde (PFA) for 45 min, washed 3x in phosphate buffered saline (PBS), and then cryoprotected with 10, 20, and 30% graded sucrose solutions at RT. Explants with culture membrane were embedded in Tissue-Tek O.C.T. compound (Sakura Finetek Europe, Alphen aan den Rijn, Netherlands), snap-frozen on liquid N_2_ and cryosectioned (12 μm thickness) on a Thermo Scientific NX50 microtome (Thermo Fisher Scientific, Waltham, MA, USA). Retinal sections were thaw mounted onto Superfrost Plus object slides (R. Langenbrinck, Emmendingen, Germany) and stored at -20°C for further use. To prepare unfixed tissue, retinal explants were directly embedded in Tissue-Tek O.C.T. (Sakura) and snap-frozen on liquid N_2_. Further processing was as above for fixed tissue.

### TUNEL assay

The terminal deoxynucleotidyl transferase dUTP nick end labeling (TUNEL) assay (Sigma-Aldrich *in situ* Cell Death Detection Kit, 11684795910, red fluorescence) was employed on retinal sections for the visualization of dying cells (Gavrieli et al., 1992). While fixed sections were rehydrated with phosphate-buffered saline (PBS; 0.1 M) for 10 min, unfixed sections were first fixed by incubation in 4% PFA for 10 min, and then washed in PBS (2x 5 min). Both types of preparations were incubated with proteinase K (1.5 µg/µL) diluted in preheated Tris-buffered saline (TBS; 1µL enzyme in 7mL TBS) at 37°C for 5 min. This was followed by incubation with ethanol acetic acid (70:30) at -20°C for 5 min, and washing with PBS 3 times for 5 min each. Subsequently, sections were incubated with blocking solution (10% normal goat serum; NGS, 1% bovine serum albumin; BSA, 1% fish gelatin in PBS, 0.3% Triton X-100) for 1h at RT. TUNEL reaction solution was then applied with overnight incubation at 4°C. Afterwards, the tissue was washed in PBS (3x 5 min) and mounted using Vectashield with DAPI (Vector Laboratories, Burlingame, CA, USA). Here, DAPI served as nuclear counterstain.

### PARP *in situ* activity assay

To visualize PARP activity in individual photoreceptor cells *in situ* (Belhadj et al., 2021), unfixed tissue sections were dried for 30 min at 37°C, rehydrated with Tris Buffer (50 mM) for 10 min, and then incubated for 3.5 h at 37 °C with PARP reaction mixture containing 1 mM dithiothreitol, 50 μM 6-Fluo-10-NAD^+^ (N023; Biolog, Bremen Germany), 10 mM MgCl_2_, 100 mM TBS, and 0.2% Triton X100. The sections were mounted in Vectashield with DAPI (Vector) for microscopy.

### Calpain *in situ* activity assay

This assay was performed on unfixed tissue sections (Belhadj et al., 2022). After thawing, retinal tissue sections were dried for 30 min at 37°C and then rehydrated for 15 min in calpain reaction buffer (CRB; 5.96 g HEPES, 4.85 g KCl, 0.47 g MgCl_2_, and 0.22 g CaCl_2_ in 100 mL ddH_2_O; pH 7.2). Afterwards, sections were placed in CRB with 2 mM dithiothreitol and tBOC-Leu-Met-CMAC (25 µM; A6520; Thermo Fisher Scientific) for 3.5 h at 37°C. After washing with PBS (2x 10 min), the sections were incubated with ToPro-3 (Thermo Fisher Scientific) as nuclear counterstain (1:1000 in PBS), for 25 min. Finally, sections were mounted using Vectashield without DAPI (Vector Laboratories Inc., Burlingame, CA, USA) for immediate visualization by microscopy.

### Immunostaining

Retinal sections were rehydrated with PBS for 15 min, then incubated in blocking solution (10% NGS, 1% BSA, and 0.3% PBST (PBS + 0.3% Triton X-100)) for 1 h at RT. This was followed by incubation with primary antibody: rabbit anti-calpain-2 (1:200; AB39165, Abcam, Cambridge, UK), rabbit anti-cone arrestin-3 (1:500; AB15282; Merck KGaA, Darmstadt, Germany), rabbit cleaved caspase-3 (1:100; D175; Cell Signaling, Danvers, USA), mouse anti-rhodopsin (1:500; MAB5316; Merck) at 4°C overnight. Afterwards, Alexa Fluor 488 anti-rabbit or 568 anti-mouse secondary antibodies (Thermo Fisher Scientific) were applied for 1 h at RT. Then the tissue was washed with PBS 3 x 5 min, followed by mounting in Vectashield with DAPI (Vector Laboratories Inc., Burlingame, CA, USA).

### Microscopy

Light and fluorescence microscopy were performed on a Zeiss Imager Z.2 microscope equipped with ApoTome 2, an Axiocam 506 mono camera, and HXP-120V fluorescent lamp (Carl Zeiss Microscopy, Oberkochen, Germany) at RT. The excitation (λExc.)/emission (λEm.) characteristics of the filter sets used for the different fluorophores were as follows (in nm): DAPI (λExc. = 369 nm, λEm = 465 nm), AF488 (λExc. = 490 nm, λEm = 525 nm), and ToPro3 (λExc. = 642 nm, λEm = 661 nm). Representative images were captured from the retina using a 20x/0.8 objective lens and Zen 2.3 Blue Edition software. Sections of 12 µm thickness were analyzed using a 9-optical section ApoTome2 Z-stack.

### Quantification and statistical analysis

Pictures were captured on three entire sagittal sections from at least three different animals. The average area occupied by a photoreceptor cell (*i*.*e*., cell size) was determined by counting DAPI-, or ToPro-3-stained nuclei in nine different areas of the retinal outer nuclear layer (ONL) to establish an average cell size. The number of positive-labeled cells in the ONL was counted manually. Arrestin-3-labeled cells in the ONL were counted manually, and their numbers were represented as cells per 100 μm of retinal circumference.

To calculate the minimum number of animals and samples required for the statistical analyses, a power calculation based on previously observed effect sizes (Sahaboglu et al., 2016; Zhu, Kumar, et al., 2025; Zhu, Peiroten, et al., 2025) was performed using the online tool StatistikGuru, Version 1.96 (www.statistikguru.de; information retrieved August 2025). Animals were not assigned to experimental groups prior to their sacrifice, both retinal explants generated from experimental animals were included in the analysis and always assigned to different experimental groups. Errors in graphs and text are given as standard deviation (SD). Values are given as mean ± SD. Statistical analysis was performed using GraphPad Prism 10.2.3 software (GraphPad Software, La Jolla, CA, USA), one-way and two-way ANOVA test with Dunnett’s multiple comparisons test were performed to compare more than two groups. Levels of significance were: ^*^ = *p* ≤ 0.05, ^**^ = *p* ≤ 0.01, ^***^ = *p* ≤ 0.001, ^****^ = *p* ≤ 0.0001. Full statistical reports are presented in Tables S1-8.

## Results

### The PARP inhibitor olaparib increases *Rho*^**I255del**^ **photoreceptor cell death**

Previous studies found that treatment with olaparib (Ola) – a PARP inhibitor that favors PARP-2 over PARP-1 (Table 1) – protected photoreceptors in the *rd1* and *rd10* mouse models for recessive RP (Dong et al., 2023; Sahaboglu et al., 2016; Vidal-Gil et al., 2019; Yan et al., 2022), but its effect on *Rho*^I255del^ mutant retina was unclear. To address this question, we employed organotypic retinal explants derived from heterozygous *Rho*^I255del/+^ (*R*^I255d/+^) and wild-type (WT) mice, cultured from post-natal (P) day 12 to P20. After two days without treatment, at P14, the explants were exposed to rising concentrations of Ola (0.1, 1, and 10 µM). Since DMSO was used as a solvent for Ola, we used DMSO-treated cultures for comparison (0.01, 0.1, and 1 %) (Tsai et al., 2009). Readouts consisted of PARP *in situ* activity on unfixed retinal sections, and TUNEL assay on 4% PFA-fixed retinal sections (Fig. 2, Table S1).

**Figure 2.**
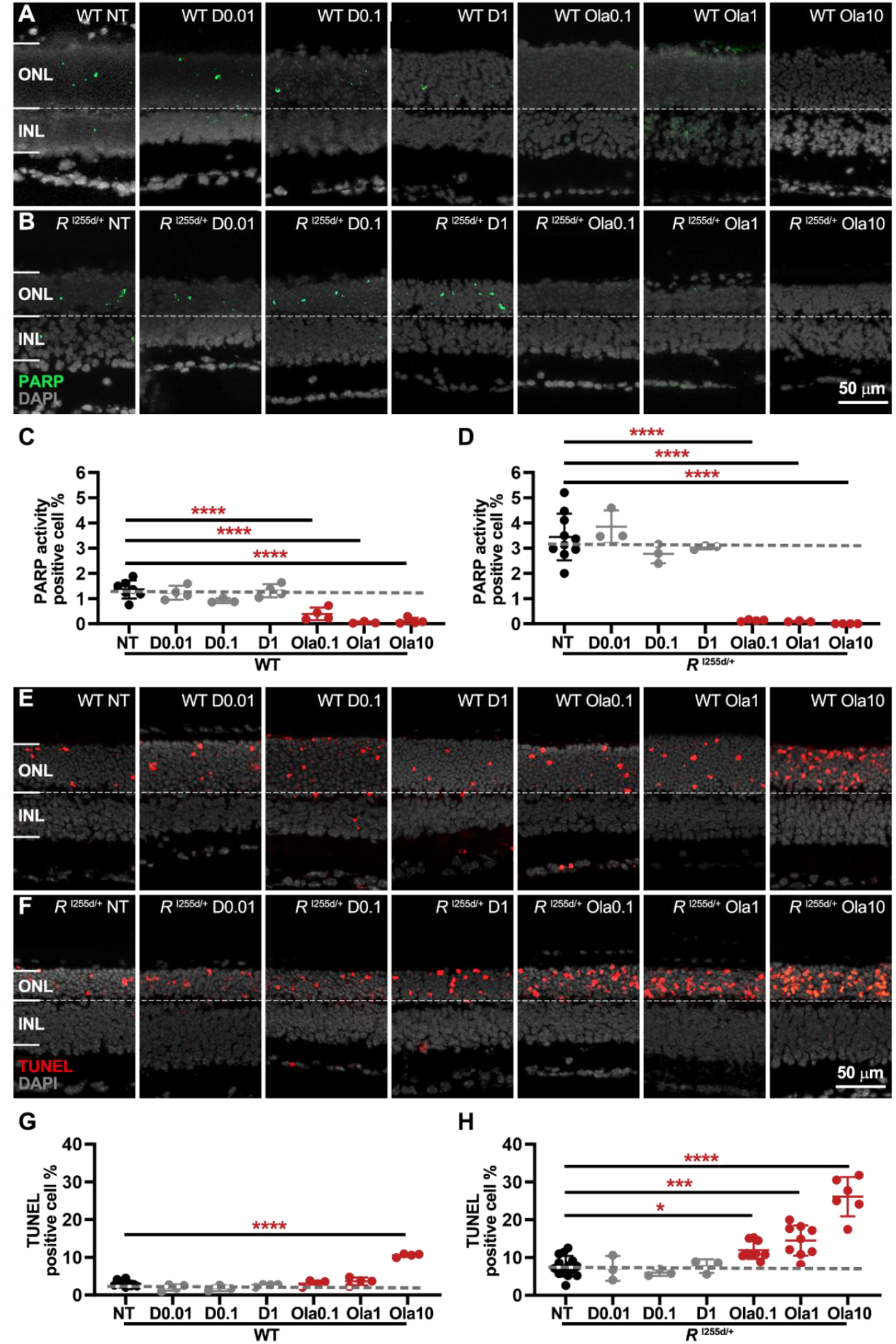
Olaparib dose-dependently reduces PARP activity and causes photoreceptor cell death. Organotypic retinal explants were derived from wild-type (WT) and *Rho*^I255d/+^ (*R*^I255d/+^) mice, and cultured from post-natal (P) day 12 to 20. Cultures were either non-treated (NT) or treated with different concentrations of DMSO (D; 0.01, 0.1, and 1%) or olaparib (Ola; 0.1, 1, and 10 µM). (**A, B**) PARP *in situ* activity assay (green) labeled cells mostly in the outer nuclear layer (ONL). (**C, D**) DMSO treatment did not decrease PARP activity, neither in WT nor in *Rho*^I255d/+^ retina. In contrast, Ola significantly reduced PARP activity in both genotypes. (**E, F**) TUNEL assay staining dying cells (red) in the ONL of both WT and *Rho*^I255d/+^. (**G, H**) Quantification of TUNEL-positive cells showed no DMSO treatment effect in WT and *Rho*^I255d/+^ retina, while Ola treatment dose-dependently increased cell death. DAPI (grey) was used as nuclear counterstain. Images represent 3-10 independent retinal explant cultures; error bars indicate SD; statistical analysis: One-way ANOVA with Dunnett’s multiple comparisons test; ^*^ = *p* ≤ 0.05; ^**^ = *p* ≤ 0.01; ^***^ = *p* ≤ 0.001; ^****^ = *p* ≤ 0.0001; INL= inner nuclear layer.

In both WT and mutant retina, exposure to DMSO did not alter PARP activity in the outer nuclear layer (ONL), when compared to non-treated (NT) retina. In contrast, Ola strongly reduced PARP activity in both WT and *Rho*^I255del/+^ retinas (Fig. 2A-D), indicating that it had inhibited PARP effectively.

The numbers of dying ONL cells, as detected by the TUNEL assay, were not changed by DMSO. Remarkably, the numbers of TUNEL-positive cells in the ONL were dose-dependently increased by Ola treatment in mutant retina and – to a lesser extent – in WT retina, when compared to NT (Fig. 2E-H). At 10 µM concentration Ola was clearly toxic to both genotypes. Since Ola targets PARP-1 and -2, these results indicate that in *Rho*^I255del/+^ retina these PARP isoforms may be neuroprotective factors.

### Olaparib increases caspase-3 activation in rod photoreceptors

Cell death is typically associated with increased proteolysis. In ARRP and ADRP animal models, retinal degeneration has previously been connected to excessive activation of Ca^2+^-dependent, calpain-type proteases and especially activation of the calpain-2 isoform (Arango-Gonzalez et al., 2014; Belhadj et al., 2022; Power et al., 2020). Increased calpain activity has also been found in *Rho*^I255d/+^ retina at P20 (Bowen Cao et al., 2024), and this activity may be related to non-apoptotic cell death, triggered by high cGMP (Power et al., 2020). In contrast, apoptotic cell death is typically connected to caspase-type proteases, in particular to activation of caspase-3 (Arango-Gonzalez et al., 2014).

To determine the effect of Ola on calpain activity, as well as on calpain-2 and caspase-3 activation, we subjected Ola-treated *Rho*^I255d/+^ retina to corresponding staining procedures (Fig. 3, Table S2). In P20 cultured *Rho*^I255d/+^ retina the numbers of cells displaying high calpain activity were not significantly changed in the ONL of NT mutant compared to WT. Yet, calpain activity was significantly increased at a concentration of 10 µM olaparib, compared to NT mutant (Fig. 3A, B). However, calpain-2 activation, which was increased in NT mutant compared to NT WT, was not altered by Ola treatment in *Rho*^I255d/+^ retina (Fig. 3C, D). Cells showing activation of caspase-3 were essentially absent from NT WT, but significantly increased in NT *Rho*^I255d/+^ ONL. In the mutant ONL, caspase-3 activation increased with Ola concentration (Fig. 3E, F), indicating that Ola treatment may have promoted the execution of apoptotic cell death.

**Figure 3.**
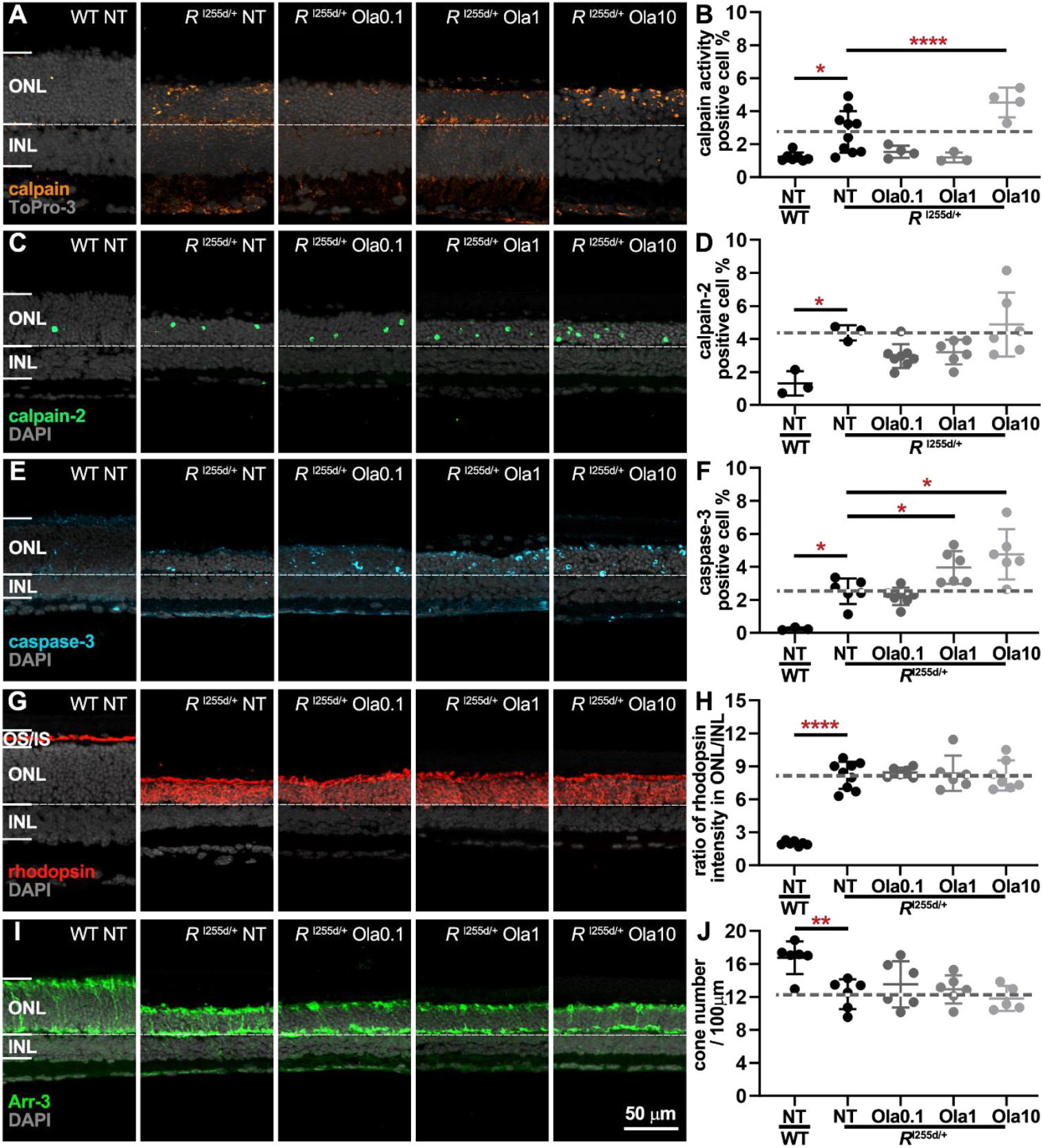
Olaparib alters proteolytic activities in *Rho*^I255d/+^ retina, without affecting cone photoreceptors. Organotypic retinal explants were derived from wild-type (WT) and *Rho*^I255d/+^ (*R*^I255d/+^) mice and cultured from post-natal (P) days 12 to 20. Cultures were either non-treated (NT) or treated with 0.1, 1, or 10 µM olaparib (Ola). Calpain *in situ* activity assay was performed on unfixed tissue sections, while immunostaining for activated calpain-2, caspase-3, rhodopsin, and arrestin-3 (Arr-3) was performed on fixed sections. (**A-B**) Calpain activity assay (orange) labeled cells in the outer nuclear layer (ONL), and their numbers were significantly increased at 10 µM Ola. (**C-D**) Compared to WT, calpain-2 activation (green) was prominently increased in the mutant retina, however, Ola treatment did not significantly change calpain-2. (**E-F**) Immunostaining for cleaved caspase-3 (cyan) produced a ring-shaped labeling in ONL cells, which was more prominent in the mutant compared to WT. The number of caspase-3-positive cells increased at 1 and 10 µM Ola compared to NT mutant retinas. (**G-H**) Rhodopsin immunostaining (red) revealed a mislocalization to the ONL in mutant retina, that was not seen in WT. Yet, Ola treatment did not alter the ratio of rhodopsin ONL/inner nuclear layer (INL) staining intensity. (**I-J**) Cone photoreceptors labeled by cone-arrestin-3 (Arr-3; green) showed a decrease in NT mutant compared to WT; however, Ola treatment did not affect cones. Nuclear counterstain (grey) with ToPro-3 (**A**) or DAPI (**C, E, G, I**). Images are representative for 3-10 independent retinal cultures; error bars indicate SD; statistical analysis: Two-way ANOVA with Dunnett’s multiple comparisons test; levels of significance were: ^*^ = *p* ≤ 0.05; ^**^ = *p* ≤ 0.01; ^****^ = *p* ≤ 0.0001.

Rod photoreceptors in *Rho*^I255d/+^ retina were previously found to have high levels of rhodopsin mislocalized to the ONL rather than to photoreceptor outer segments (OS) (Bowen Cao et al., 2024). To assess whether Ola had altered rhodopsin trafficking, we used immunostaining and compared the rhodopsin staining intensity between inner nuclear layer (INL) and ONL (Fig. 3G, H). The ratio of rhodopsin staining intensity ONL/INL showed a clear increase in *Rho*^I255d/+^ retina compared to WT. This ratio was not affected by rising Ola levels, suggesting that rod photoreceptor rhodopsin localization was not changed by the treatment.

Since in the mouse ONL rod photoreceptors outnumber cone photoreceptors approx. 20:1 (Carter-Dawson & LaVail, 1979) (*i*.*e*. cones constitute approx. 5% of all ONL cells), the very high cell death numbers seen with Ola treatment suggested that mostly rods were affected by the treatment. To nonetheless evaluate possible effects of Ola on cone viability, we used immunostaining for the cone marker arrestin-3 (Arr-3). Remarkably, while the number of cones was generally decreased in mutant retina compared to WT, Ola did not alter cone survival in mutant retina. Overall, these results suggest that Ola-mediated PARP inhibition had a strong detrimental effect on rod but not cone photoreceptors. This detrimental effect appeared to be connected to increased caspase-3-dependent apoptosis but seemed independent of mislocalized rhodopsin.

### Destructive effects of olaparib do not depend on culture duration

The negative effect of Ola on photoreceptor degeneration at P20, while showing strong PARP inhibition, was unexpected. To investigate whether this effect was somehow influenced by the duration of the treatment, we decreased the culturing period by two days to end at P18, *i*.*e*. the earliest time-point at which the *Rho*^I255del/+^ mutant displays a significant increase in PARP activity and photoreceptor cell death (Bowen Cao et al., 2024). Also at P18, 1 µM Ola treatment strongly reduced PARP activity in mutant retina (Fig. S1A-B, Table S3-4), while not affecting ONL calpain activity (Fig. S1C-D). The TUNEL assay confirmed a strong increase in ONL cell death under Ola treatment (Fig. S1E-F). These results indicated that PARP inhibition triggered *Rho*^I255d/+^ photoreceptor cell death already at P18, *i*.*e*. prior to the peak of cell death in the *Rho*^I255del/+^ mutant retina. To facilitate a comparison of Ola effects in short (P18) and long term (P20) treatment, we plotted PARP activity, calpain activity, cell death, and ONL row counts against culture duration (Fig. S1G-J). There was a tendency for all four parameters to decrease from P18 to P20, reflecting the progression of the degeneration in this time period.

### The PARP-1 inhibitor saruparib causes photoreceptor cell death in *Rho*^**I255d/+**^ **retina**

While Ola inhibits both PARP-1 and -2 isoforms, saruparib (Sar) is a highly selective PARP-1 inhibitor (Table 1). To assess whether the detrimental effects observed with olaparib were due to PARP-1 or PARP-2 inhibition, we exposed *Rho*^I255d/+^ organotypic retinal explant cultures to Sar, using the same experimental paradigm as for Ola.

With Sar treatment, the numbers of *Rho*^I255d/+^ ONL cells displaying high PARP activity decreased in a dose dependent fashion (Fig. 4A-B, Table S5). Conversely, in mutant retinas calpain activity-positive cells increased with rising Sar concentration, compared to the NT situation (Fig. 4C-D). This increase in calpain activity, already at low Sar concentrations, was not seen with Ola treatment (*cf*. Fig. 3B). *Rho*^I255d/+^ ONL cell death (TUNEL assay) showed a similar Sar dose-dependent increase (Fig. 4E-F). These results suggest that some of the PARP activity seen in the *Rho*^I255d/+^ ONL likely originates from PARP-2, and that specific PARP-1 inhibition induces photoreceptor cell death and calpain activity in the *Rho*^I255d/+^ retina.

**Figure 4.**
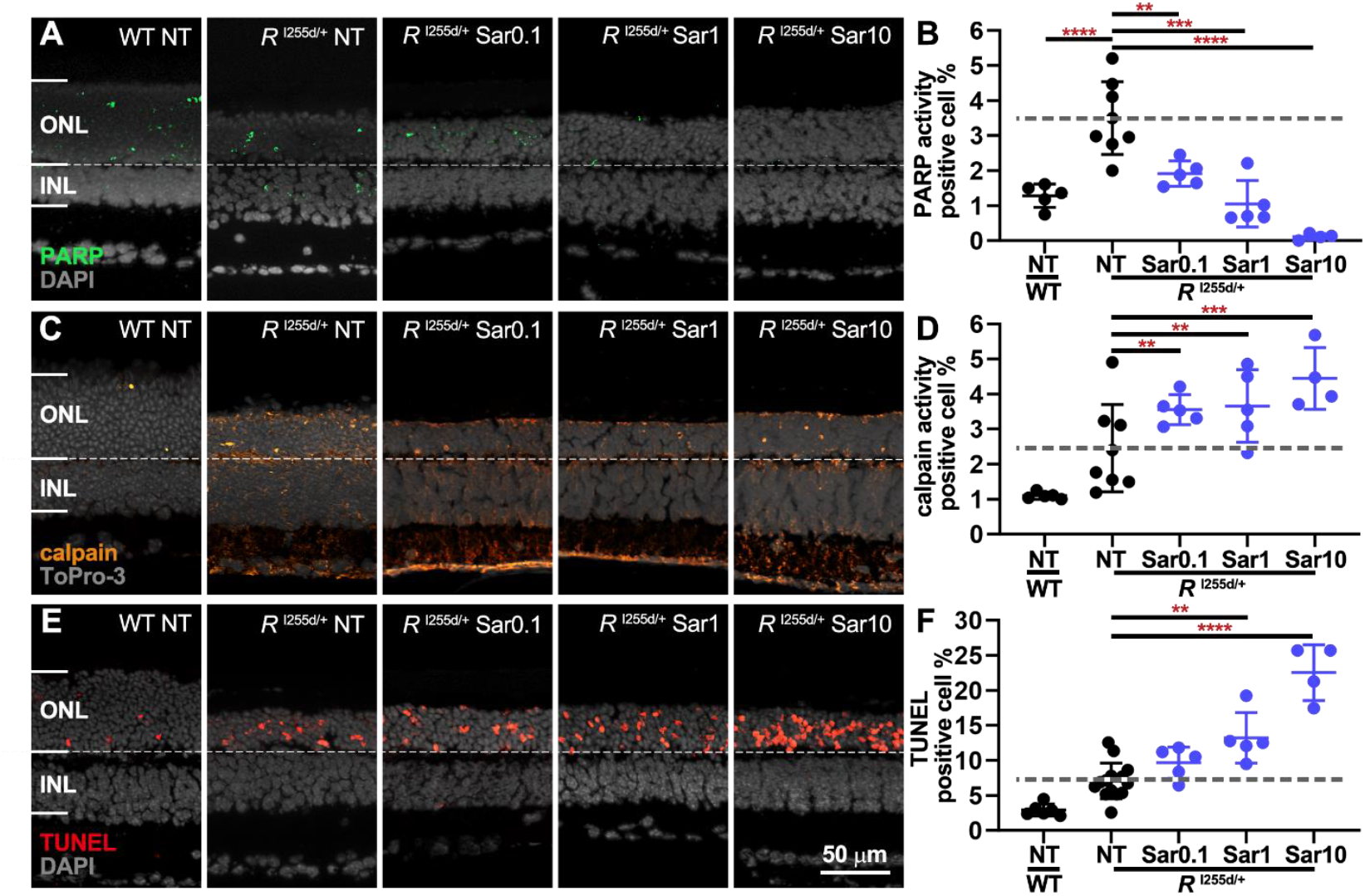
Saruparib decreases PARP activity; increases calpain activity and photoreceptor cell death. Organotypic retinal explants were derived from *Rho*^I255del/+^ (*R*^I255d/+^) mice and cultured from post-natal (P) day 12 to 20. Cultures were either non-treated (NT) or treated with 0.1, 1, or 10 µM saruparib (Sar). TUNEL-assay, PARP- and calpain-*in situ* activity were used as readouts. Wild-type (WT) data shown for comparison. (**A-B**) Sar treatment significantly reduced PARP-positive cells (green) in the outer nuclear layer (ONL). (**C-D**) Calpain activity-positive cells (orange) were increased by Sar treatment. (**E-F**) ONL cell death, visualized via TUNEL assay (red), was dose-dependently increased by Sar treatment. DAPI (**A, E**) and ToPro-3 (**C**) served as nuclear counterstains (grey). Images are representative for 4-14 independent retinal cultures; error bars indicate SD; statistical analysis: Two-way ANOVA with Dunnett’s multiple comparisons test; levels of significance were: ^**^ = *p* ≤ 0.01; ^***^ = *p* ≤ 0.001; ^****^ = *p* ≤ 0.0001; INL= inner nuclear layer.

### INO1001 inhibits PARP activity and triggers photoreceptor cell death in *Rho*^I255d/+^ retina at high doses

To confirm the detrimental effects of PARP inhibition, we used a third compound, INO1001 (INO), which displays a marked (30-fold) preference for PARP-1 over PARP-2 (Pires et al., 2025)(Table 1). Using the same test paradigm as before, we found INO to have a relatively weak effect on the numbers of PARP-activity positive cells in the *Rho*^I255d/+^ ONL, an effect that was significant only at 10 µM concentration (Fig. 5A-B, Table S6). As opposed to the treatments with Ola and Sar, INO did not change calpain activity in the mutant ONL (Fig. 5C-D). Nevertheless, also INO exhibited a clear detrimental effect on *Rho*^I255d/+^ photoreceptor viability, as indicated by the increase in the number of TUNEL-positive cells (Fig. 5E-F). To sum up, INO inhibited PARP activity and induced photoreceptor cell death at high concentration.

**Figure 5.**
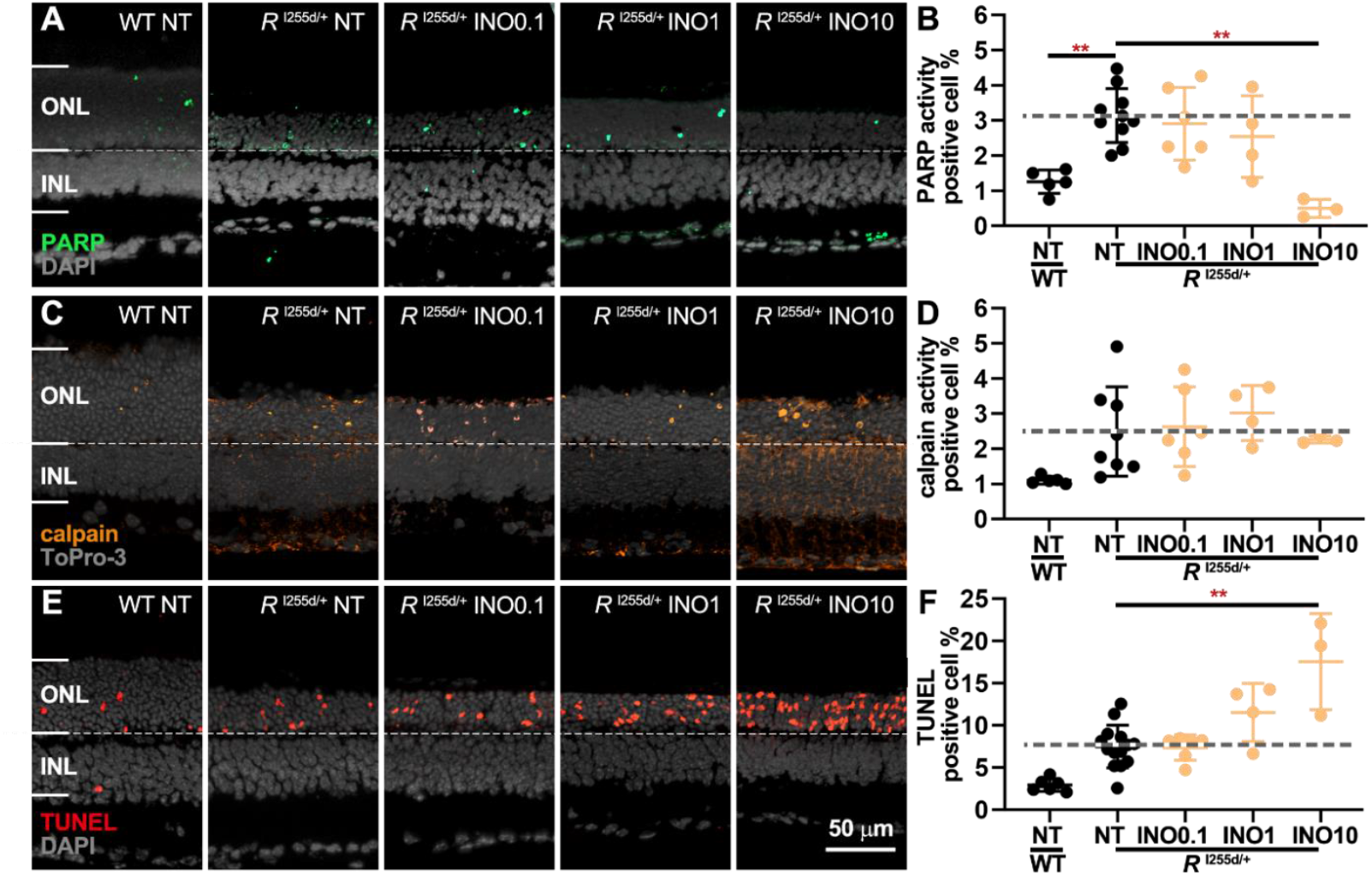
INO1001 inhibits PARP activity and induces photoreceptor cell death at highest concentration. Organotypic retinal explants were derived from *Rho*^I255del/+^ (*R*^I255d/+^) mice and cultured from post-natal (P) day 12 to 20. Cultures were either non-treated (NT) or treated with 0.1, 1, or 10 µM INO1001 (INO). PARP- and calpain-*in situ* activity and TUNEL assays were used as readouts. Wild-type (WT) data shown for comparison. (**A-B**) The numbers of PARP-activity positive cells (green) in the outer nuclear layer (ONL) of the mutant retina were significantly reduced by 10 µM INO treatment. (**C-D**) The numbers of calpain activity assay positive cells (orange) in the ONL were not changed by INO treatment. (**E-F**) ONL cell death (TUNEL assay, red) was significantly increased by 10 µM INO treatment. DAPI (**A, E**) and ToPro-3 (**C**) served as nuclear counterstains (grey). Images are representative for 3-14 independent retinal cultures; error bars indicate SD; statistical analysis: Two-way ANOVA with Dunnett’s multiple comparisons test; levels of significance were: ^**^ = *p* ≤ 0.01; INL= inner nuclear layer.

### Nicotinamide decreases PARP activity, displays *Rho*^I255d/+^ photoreceptor toxicity

The PARP inhibitors Ola, Sar, and INO all exhibit some degree of DNA-trapping, *i*.*e*. they lock the PARP protein onto the DNA, impeding gene transcription or repair of DNA-damage (Murai et al., 2012; Pires et al., 2025). To ascertain that the toxicity of PARP inhibition was not caused by DNA-trapping and to ultimately settle the negative effects of PARP inhibition on photoreceptor viability in the *Rho*^I255del/+^ mutant situation, we employed nicotinamide (NAM) as a classical inhibitor of PARP activity (Clark et al., 1971), that only blocks catalytic activity but does not cause DNA-trapping. Using the above established test paradigm, we found NAM to also reduce PARP activity in *Rho*^I255d/+^ ONL (Fig. 6A, B, Table S7), albeit at much higher concentrations than the other inhibitors used in this study, in line with NAM’s relatively high IC_50_ value (*cf*. Table 1). While calpain activity was not changed by NAM treatment (Fig. 6C, D), the numbers of dying cells in the ONL were markedly and highly significantly increased at NAM concentrations of 200 µM or higher (Fig. 6E, F). This data conclusively showed that in *Rho*^I255d/+^ retina PARP inhibition was strongly detrimental to rod photoreceptors.

**Figure 6.**
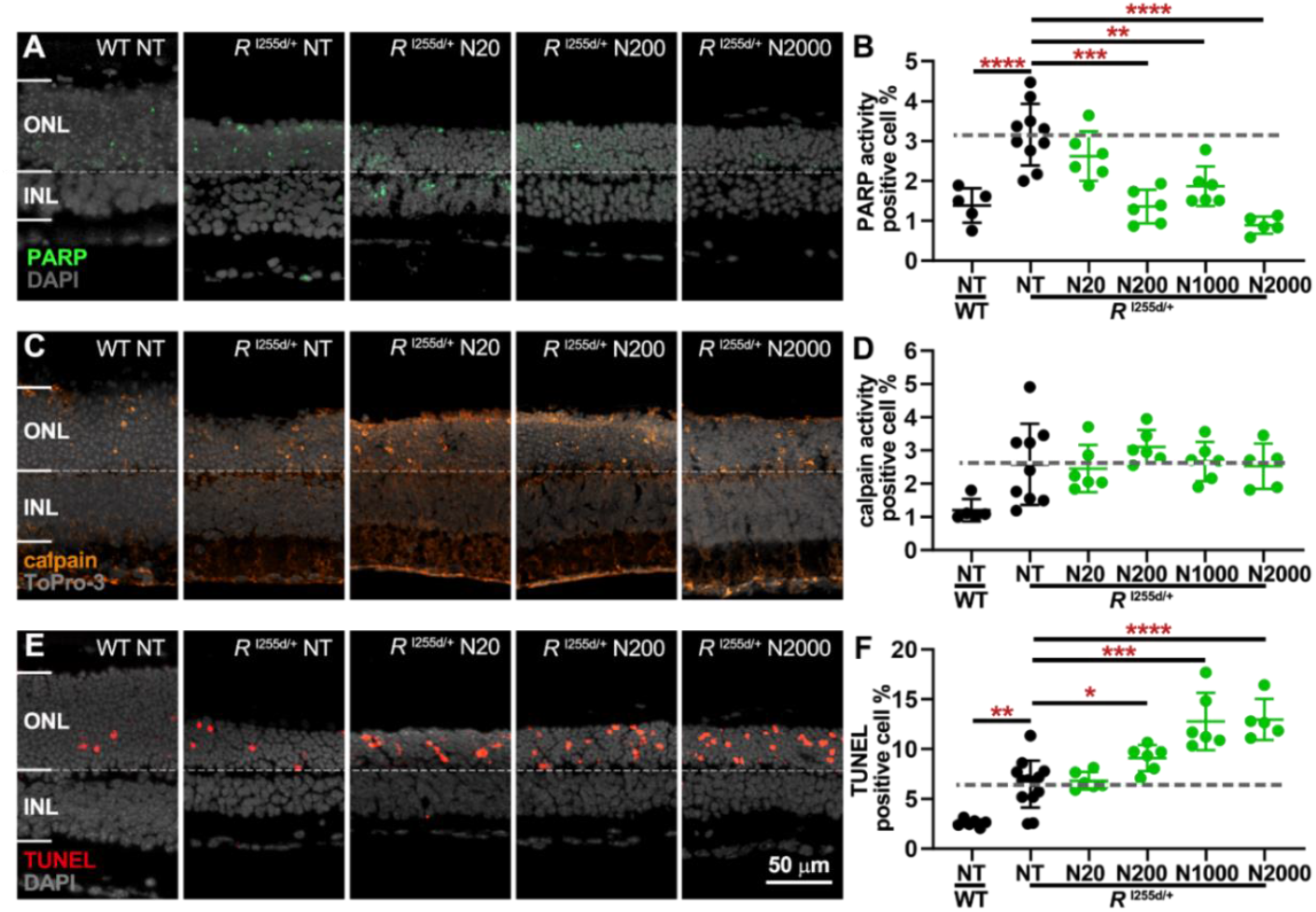
NAM decreases PARP activity, increases photoreceptor death. Organotypic retinal explants were derived from *Rho*^I255d/+^ mice at post-natal (P) day 12 and cultured with 20, 200, 1000, or 2000 µM of nicotinamide (NAM; N) from P14 to P20. Wild-type (WT) data shown for comparison. (**A, B**) In mutant retina, PARP activity (green) was dose-dependently decreased by NAM treatment. (**C, D**) Calpain activity (orange) in *Rho*^I255d/+^ ONL was not changed by NAM. (**E, F**) ONL cell death (TUNEL assay, red) was dose-dependently increased by NAM treatment. DAPI (**A, E**) and ToPro-3 (**C**) were used as nuclear counterstains (grey). Data represent mean and SD, and were obtained from 5-13 independent retinal explant cultures; statistical analysis: Two-way ANOVA with Dunnett’s multiple comparisons test; ^*^ = *p* ≤ 0.05; ^**^ = *p* ≤ 0.01; ^***^ = *p* ≤ 0.001; ^****^ = *p* ≤ 0.0001; INL= inner nuclear layer.

### Olaparib shows the strongest detrimental effects on *Rho*^I255d/+^ retina

To compare the effects of the four PARP inhibitors tested – olaparib (Ola), saruparib (Sar), INO1001 (INO), and nicotinamide (NAM) (Table 1) – we plotted the combined data on PARP activity, calpain activity, cell death and ONL row counts in summary graphs (Fig. 7, Table S8). This comparison revealed that Ola had the strongest effect on PARP activity in the *Rho*^I255d/+^ ONL and produced the most marked increase in ONL cell death. Conversely, NAM had the overall weakest effect on these parameters. Sar caused a strong rise in calpain activity already at relatively low concentrations, an effect that seemed disproportional to PARP inhibition, and which could indicate further off-target effects of Sar. The number of ONL rows were not significantly affected by any of the PARP inhibitors used here, within the time-frame of the experiment. Yet, it seems likely that the increased cell death rates would have translated into stronger loss of ONL row counts, if culturing had been prolonged by a few more days.

**Figure 7.**
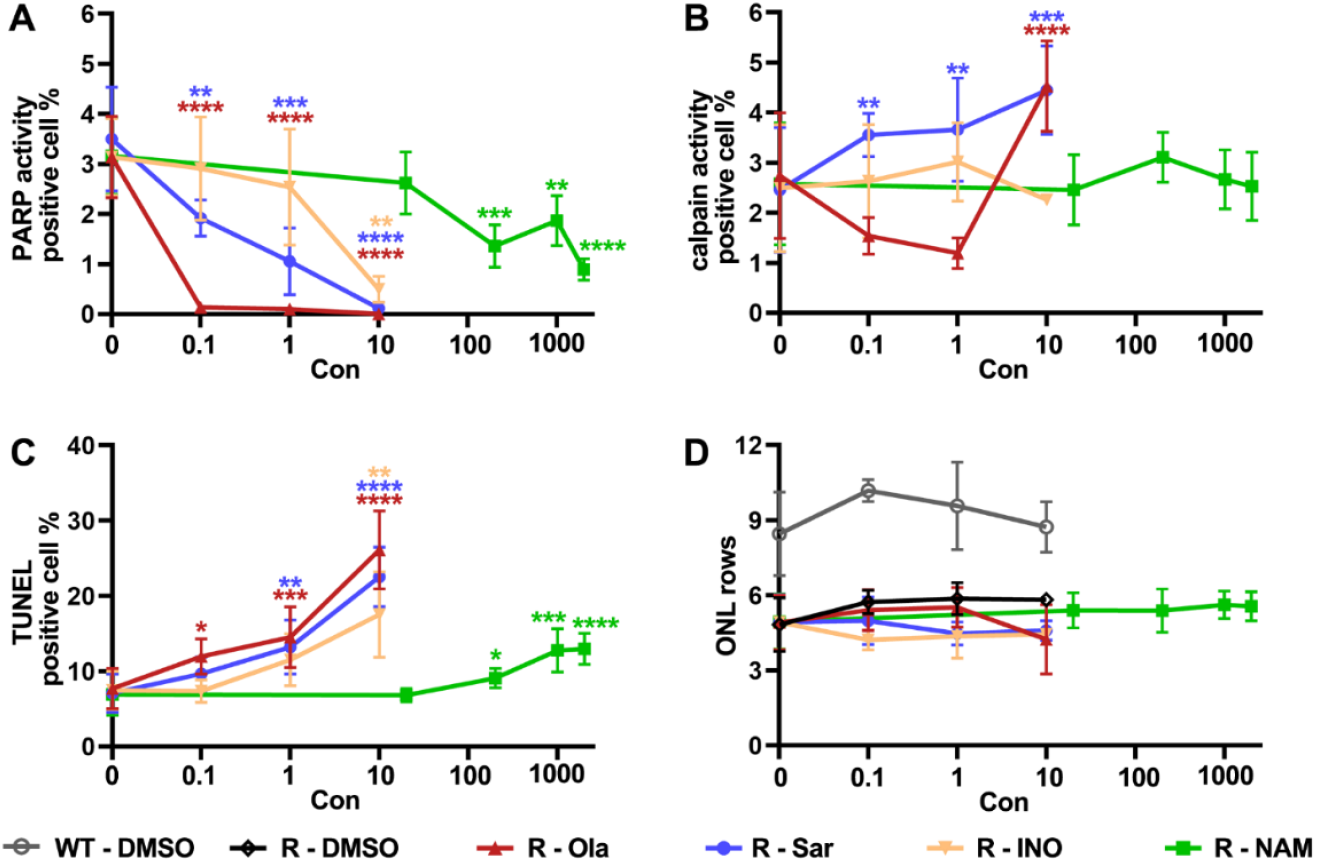
Inhibitors dose-dependently reduce PARP activity and increase *Rho*^*I255d*/+^ photoreceptor death. Summary graphs for the effects of treatment with olaparib (Ola, dark red), saruparib (Sar, blue), INO1001 (INO, light brown), and nicotinamide (NAM, green). DMSO (black/grey) shown as control. (**A**) The number of ONL cells displaying high PARP activity were reduced by all PARP inhibitors in a dose-dependent manner, with Ola showing the strongest and NAM the weakest effect. (**B**) ONL calpain activity was increased by Sar treatment. (**C**) All four PARP inhibitors dose-dependently increased ONL cell death. (**D**) In WT retina and *Rho*^*I255del*/+^ retina, DMSO did not significantly change the ONL cell row counts. In mutant retina PARP ONL row counts were not affected by PARP inhibitors within the experimental time frame (post-natal day 20). Error bars indicate SD; statistical analysis: Two-way ANOVA with Dunnett’s multiple comparisons test; ^*^ = *p* ≤ 0.05; ^**^ = *p* ≤ 0.01; ^***^ = *p* ≤ 0.001; ^****^ = *p* ≤ 0.0001.

## Discussion

Elevated PARP activity has been found in degenerating photoreceptors in a high number of animal models for RP and other inherited retinal degenerations (Arango-Gonzalez et al., 2014), including in the recently generated *Rho*^I255d^ ADRP mouse (Bowen Cao et al., 2024). This finding suggested using over-activated PARP as a common target for therapeutic interventions in a large group of highly heterogenic genetic diseases (Verbakel et al., 2018). Our present study however provides evidence that in the photoreceptors of *Rho*^I255d/+^ autosomal-dominant rhodopsin mutants, PARP activity is beneficial and required, so that its inhibition rather aggravates the disease phenotype. This highlights the need for a more comprehensive insight into underlying disease mechanisms to guide effective and safe future therapy developments.

### A novel, apparently ambiguous role of PARP in neurodegenerative diseases

A property of PARP enzymes is to sense DNA damage, which triggers poly(ADP-ribosyl)ation, a process in which PARP transfers ADP-ribose moieties from NAD^+^ to target proteins, like the histones of the chromatin (Curtin & Szabo, 2020). A such activity is crucial for DNA repair, PARP has been referred to as the “guardian of the genome” (Chatterjee et al., 1999). The protective role of PARP was the basis for the development of PARP inhibitors as anti-cancer drugs, which, especially when administered in combination with DNA-damaging compounds or radiation, promote the death of mitotically active cells (Bhamidipati et al., 2023). This contrasts with excessive PARP activation frequently observed in neurodegenerative diseases, which has led to the concept of PARP-mediated cell death or PARthanatos (Yang et al., 2024). Accordingly, PARP inhibitors originally developed for cancer therapy may serve as possible neuroprotective agents. In murine models for autosomal-recessive RP of the retina, carrying homozygous disease-causing mutations in *Pde6a* and *Pde6b* genes, inhibition of PARP with *e*.*g*. PJ34 or olaparib was found to significantly delay photoreceptor degeneration (Dong et al., 2023; Jiao et al., 2016; Paquet-Durand et al., 2007; Sahaboglu et al., 2016; Yan et al., 2022). The protective effect of PARP inhibition has been attributed to a decreased consumption of NAD^+^, which is a critical cofactor in ATP-producing energy metabolism. According to this hypothesis, overactivation of PARP may severely impair the capacity of a cell to produce ATP (Du et al., 2003), while PARP inhibition in contrast would allow to maintain cellular ATP at essential levels.

The work presented here obtained in the *Rho*^I255d/+^ model of autosomal-dominant RP, carrying one normal and one mutated allele, is in contradiction with earlier results derived from homozygous autosomal-recessive RP models. The cell death-promoting effects of the four different PARP inhibitors used in the present study unmistakably demonstrate that in *Rho*^I255d/+^ photoreceptors PARP activity assumes a protective rather than a destructive role. As the main difference between autosomal dominant and autosomal recessive RP is the presence of a regular, unaffected allele in the further, a reason for the findings may be that PARP inhibition may have differential effects on gene activation, transcription, and protein translation between diseased and normal alleles. Alternatively, the damaging effects of PARP inhibitors in a rhodopsin mutant may relate to the relatively high transcription rate of the rhodopsin gene (DesJardin et al., 1993), since PARP activity is required for the initiation of gene transcription (Kraus & Hottiger, 2013). Still, we did not observe changes in photoreceptor rhodopsin expression or (mis-)localization under olaparib treatment. As in all autosomal-dominant forms of RP, the I255del mutation is expected to lead to misfolding and aggregation of irregular rhodopsin protein in the endoplasmic reticulum (ER), and the ensuing unfolded protein response (UPR) may consume large amounts of ATP (B. Cao et al., 2024) that would not be correctable by PARP inhibition. Rather, a sufficient degree of PARP activity may be required for the proper degradation of misfolded rhodopsin, since at least one PARP isoform – PARP16 – was found to be a key regulator of UPR (Jwa & Chang, 2012). Thus, it is plausible to think that PARP – or specific PARP isoforms – may be required to uphold adequate UPR function, and that in a condition of strong unfolded/misfolded protein overload an inhibition of PARP is detrimental. This hypothesis would explain why PARP inhibition is beneficial in autosomal-recessive RP forms where protein misfolding and UPR are usually not critical, and where reduced consumption of NAD^+^ and reduced strain on energy metabolism would improve cellular survival. Furthermore, *Rho*^I255d/+^ cone photoreceptors would not be affected by a possible (mis-)regulation of UPR by PARP inhibitors due to different opsin types, expression rates, and transport mechanisms.

### PARP inhibition: Switching from non-apoptotic to apoptotic cell death?

Photoreceptor loss is strongly linked to (excessive) PARP activity (Belhadj et al., 2021; Paquet-Durand et al., 2007), however, the exact underlying cell death mechanisms and how PARP may be involved in these is still unclear. Other than to PARthanatos, excessive activation of PARP may be connected to further, alternative cell death pathways, including necroptosis and cGMP-dependent cell death (Bowen Cao et al., 2024; Power et al., 2020; Virág et al., 2013; Zhu et al., 2022). In animal models for both ARRP and ADRP, including the *Rho*^I255d/+^ model (Zhu, Peiroten, et al., 2025), photoreceptor degeneration has been connected specifically to excessive cGMP-signaling and non-apoptotic forms of cell death (Power et al., 2020). In addition, PARP has also been proposed to be involved in the execution of apoptosis, although here the links are far less clear (Virág et al., 2013), notably because apoptosis as an active, ATP-dependent process (Liu et al., 1996) cannot be executed when NAD^+^ has been depleted (by PARP). In other words, the inhibition of PARP and the maintenance of cellular NAD^+^ and ATP at levels sufficiently high to allow for the execution of apoptosis, could promote apoptotic cell death. This corresponds to our observation on increased caspase-3 activation – a prototypical marker for apoptosis – in *Rho*^I255d/+^ photoreceptors treated with Ola, which is in line with earlier studies in other cellular systems (Sauriol et al., 2023; Zhao et al., 2019). Thus, the activity of PARP or the lack thereof in rod photoreceptors may be able to switch the execution of cell death from non-apoptotic to apoptotic mechanisms.

### Cross-talk between PARP and calpain

Apart from PARP, photoreceptor cell death is also strongly linked to calpain activity, a Ca^2+^-dependent protease that plays important functions under physiological and pathological conditions in the retina and other organs (Arango-Gonzalez et al., 2014). In degenerative retinal diseases high calpain activity has been attributed to mutation-dependent Ca^2+^-influx, failures of energy metabolism, and lack of ATP-dependent Ca^2+^-extrusion (Das et al., 2022). In the *rd1* mouse model, inhibition of PARP blocked calpain activity, while inversely inhibition of calpain did not reduce PARP activity, suggesting that PARP activity was upstream of calpain (Yan et al., 2022). In the present study, we thus expected to see a progressive decrease in calpain activity with rising PARP inhibitor concentrations. However, what we observed instead was either no change or – when treated with Sar or high-dose Ola – an increase in calpain activity.

While additional work is needed to explore this phenomenon further, we reason that in *Rho*^I255d/+^ photoreceptors, under conditions of misfolded protein overload and a block of UPR by PARP inhibitors, cells may invoke other proteolytic systems. In this context, we note that the ER is an important store for intracellular Ca^2+^, and a dysfunction induced by PARP inhibitors could lead to excessive Ca^2+^ release and subsequent calpain activation. Prolonged release of Ca^2+^ from the ER may also disrupt mitochondrial membrane potential (Tripathi & Chaube, 2012), which would lead to ATP-depletion as well as to additional Ca^2+^ release. Both processes would promote calpain activation even though release from intracellular Ca^2+^ may not be strong enough to attain the micromolar Ca^2+^-levels required to activate calpain-2. This would thus explain why under Ola treatment overall calpain activity was significantly increased while calpain-2 activation was not.

## Conclusion

In this work, we assessed the impact of different PARP inhibitors on the progression of photoreceptor degeneration in a novel, human-homologous rhodopsin-mutant *Rho*^I255del/+^ mouse model for autosomal dominant RP. So far, elevated PARP activity had been identified as a factor promoting neurodegeneration in animal models of autosomal recessive RP forms. Contrary to these studies performed mostly in *Pde6*-mutant lines, the treatment data obtained here, using four different and well-established compounds, conclusively demonstrate the destructive effects of PARP inhibition in the autosomal dominant rhodopsin-mutant *Rho*^I255del^ model. These detrimental effects may be due to the coexistence of regular and mutated alleles, a misregulation of the unfolded protein response, and/or a shift in degenerative mechanisms from cGMP-dependent pathways to classical apoptosis. The latter option may be governed by PARP-inhibition mediated improvements in photoreceptor energy metabolism. Our observations highlight the importance of carefully considering potential fields of application when utilizing PARP inhibition as a therapeutic strategy, especially in the context of retinal degeneration. Moreover, since PARP inhibitors were initially developed for anti-cancer therapy, and since adverse effects were also observed in WT retina, the widespread use of compounds like olaparib may raise concerns about potential side effects on the retina.

## Acknowledgments

The authors would like to thank Norman Rieger (Institute for Ophthalmic Research, Eberhard-Karls-Universität Tübingen) for excellent technical assistance.

## Author contributions

Conceptualization, Y.Z. and F.P.-D.; methodology, Y.Z., and A.F.; software, Y.Z.; validation, Y.Z.; formal analysis, Y.Z.; investigation, Y.Z., and A.F.; data curation, Y.Z.; writing-original draft preparation, Y.Z.; writing—review and editing, F.P.-D., M.S., and K.J.; supervision, F.P.-D.; project administration, F.P.-D.; funding acquisition, F.P.-D. All authors have read and agreed to the published version of the manuscript.

## Data Availability Statement

All data generated or analyzed during this study are included in this published article and its supplementary materials files.

## Ethics approval and consent to participate

All procedures were performed in accordance with the ARVO statement for the use of animals in ophthalmic and visual research. Animal protocols compliant with §4 of the German law of animal protection were reviewed and approved by the Tübingen University committee on animal protection (Einrichtung für Tierschutz, Tierärztlicher Dienst und Labortierkunde, Registration No. AK02/19M).

