## Supplemental figure and tables for "PARP activity is essential for retinal photoreceptor survival in the human homologous *Rho*^I255del^ mouse model for autosomal dominant retinitis pigmentosa"

### Supplemental figures

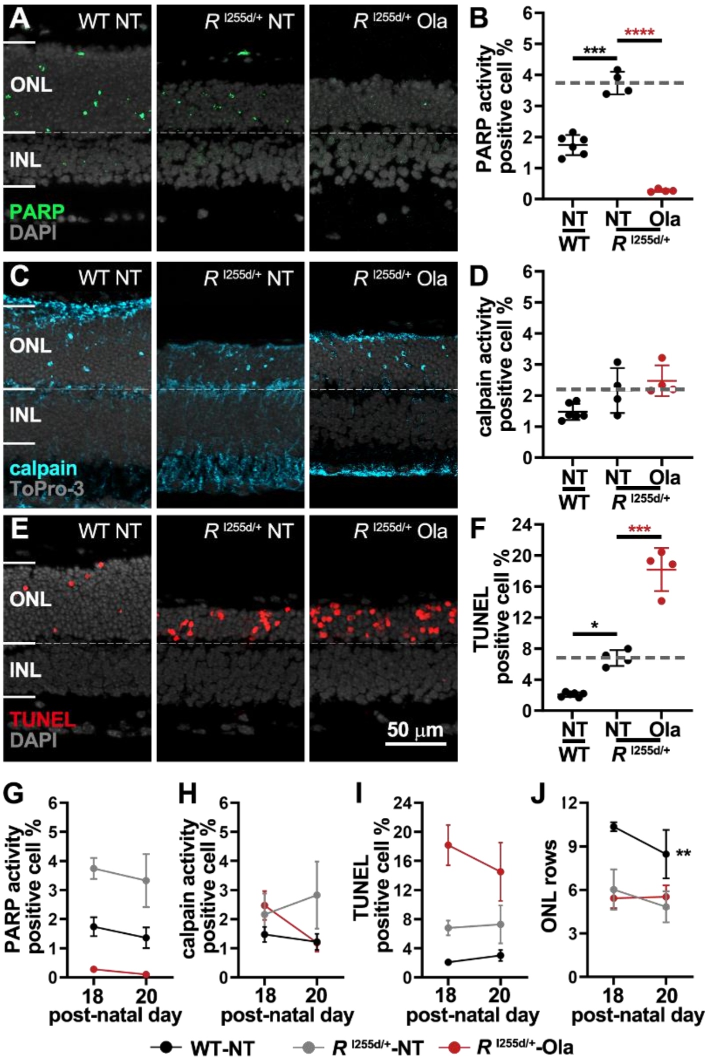


**Figure S1. Olaparib reduces PARP activity and triggers *Rho*$^{I255d/+}$ photoreceptor cell death already at P18.** Organotypic retinal explants were derived from wild-type (WT) and *Rho*$^{I255d/+}$ (*R*$^{I255d/+}$) mice and cultured from post-natal (P) day 12 to P18. Cultures were either non-treated (NT) or treated with 1 µM olaparib (Ola). (**A-B**) In P18 retina, PARP activity (green) was increased in the mutant compared to WT, but strongly decreased by Ola treatment. (**C-D**) The overall activity of calpain (cyan) in the *Rho*$^{I255d/+}$ mutant was not changed by Ola treatment. (**E-F**) Cell death in the outer nuclear layer (ONL) as indicated by the TUNEL assay (red), was significantly increased in mutant retina compared to WT. Cell death was strongly increased by Ola treatment. (**G**-**J**) Comparison of Ola treatment effects in retinal explants cultured until P18 and P20. (**G**) In both WT and mutant NT retina PARP activity decreased slightly from P18 to P20. (**H**) From P18 to P20 overall calpain activity increased in NT mutant retinas, and 1µM Ola treatment slightly reduced calpain activity in the mutant situation. (**I**) The TUNEL assay showed a minor increase in cell death in both NT WT and mutant retina from P18 to P20. In Ola-treated retina, cell death appeared to be reduced. (**G**) The number of ONL rows, as an independent marker for photoreceptor survival, decreased in NT WT. Ola treatment did not significantly improve photoreceptor survival. DAPI (**A**, **E**) and ToPro-3 (**C**) were used as nuclear counterstains (grey). Images are representative of results obtained in 3-20 independent retinal cultures; error bars indicate SD; statistical analysis: Two-way ANOVA with Dunnett's multiple comparisons test; * = $p \leqslant 0.05$; ** = $p \leqslant 0.01$; *** = $p \leqslant 0.001$; **** = $p \leqslant 0.0001$; * on the top: comparison of NT at same timepoint; * on the right: comparison of P20 with P18. INL = inner nuclear layer.

### Supplemental Tables

**Table S1.** The effect of olaparib (Ola) on PARP activity (PARP *in situ* assay) and photoreceptor cell death (TUNEL) in WT and *Rho*$^{I255d/+}$: Quantitative data for graphs presented in Figure 2 C, D, G, H.

| Table S1 (Figure) | Parameter | *p* - value comparison | Genotype-treatment | Mean ± SD (%) | *p* - value | n |
|---|---|---|---|---|---|---|
| Fig. 2C | PARP activity | WT - NT | WT - NT | 1.36 ± 0.36 | | 7 |
| | | | WT - D 0.01 | 1.23 ± 0.27 | $p = 0.9383$ | 4 |
| | | | WT - D 0.1 | 0.92 ± 0.09 | $p = 0.1042$ | 3 |
| | | | WT - D 1 | 1.31 ± 0.26 | $p = 0.9996$ | 4 |
| | | | WT - Ola 0.1 | 0.39 ± 0.25 | $p < 0.0001$ | 4 |
| | | | WT - Ola 1 | 0.05 ± 0.04 | $p < 0.0001$ | 3 |
| | | | WT - Ola 10 | 0.11 ± 0.13 | $p < 0.0001$ | 4 |
| Fig. 2D | | *R*$^{I255d/+}$ - NT | *R*$^{I255d/+}$ - NT | 3.45 ± 0.93 | | 10 |
| | | | *R*$^{I255d/+}$ - D 0.01 | 3.85 ± 0.65 | $p = 0.8792$ | 3 |
| | | | *R*$^{I255d/+}$ - D 0.1 | 2.78 ± 0.38 | $p = 0.4733$ | 3 |
| | | | *R*$^{I255d/+}$ - D 1 | 3.04 ± 0.08 | $p = 0.8723$ | 3 |
| | | | *R*$^{I255d/+}$ - Ola 0.1 | 0.13 ± 0.03 | $p < 0.0001$ | 4 |
| | | | *R*$^{I255d/+}$ - Ola 1 | 0.10 ± 0.02 | $p < 0.0001$ | 3 |
| | | | *R*$^{I255d/+}$ - Ola 10 | 0.00 ± 0.00 | $p < 0.0001$ | 4 |
| Fig. 2G | TUNEL | WT - NT | WT - NT | 3.03 ± 0.77 | | 10 |
| | | | WT - D 0.01 | 2.01 ± 0.81 | $p = 0.1498$ | 4 |
| | | | WT - D 0.1 | 1.81 ± 0.78 | $p = 0.1041$ | 3 |
| | | | WT - D 1 | 2.59 ± 0.36 | $p = 0.8792$ | 4 |
| | | | WT - Ola 0.1 | 3.10 ± 0.77 | $p > 0.9999$ | 4 |
| | | | WT - Ola 1 | 3.63 ± 1.06 | $p = 0.6479$ | 4 |
| | | | WT - Ola 10 | 10.57 ± 0.46 | $p < 0.0001$ | 4 |
| Fig. 2H | | *R*$^{I255d/+}$ - NT | *R*$^{I255d/+}$ - NT | 7.68 ± 2.70 | | 14 |
| | | | *R*$^{I255d/+}$ - D 0.01 | 7.12 ± 3.28 | $p = 0.9999$ | 3 |
| | | | *R*$^{I255d/+}$ - D 0.1 | 5.96 ± 0.86 | $p = 0.9445$ | 3 |
| | | | *R*$^{I255d/+}$ - D 1 | 7.69 ± 1.81 | $p > 0.9999$ | 3 |
| | | | *R*$^{I255d/+}$ - Ola 0.1 | 11.95 ± 2.35 | $p = 0.0232$ | 9 |
| | | | *R*$^{I255d/+}$ - Ola 1 | 14.51 ± 4.02 | $p = 0.0001$ | 9 |
| | | | *R*$^{I255d/+}$ - Ola 10 | 26.11 ± 5.20 | $p < 0.0001$ | 6 |

**Table S2.** Effect of olaparib (Ola) on calpain activity (calpain *in situ* assay), calpain-2 (calpain-2 staining) and caspase-3 activation (caspase-3 staining), rhodopsin (rhodopsin staining), and cone survival (cone arrestin-3 staining) in outer nuclear layer (ONL) or/and inner nuclear layer (INL), with concentrations (0.1, 1, and 10 µM): Quantitative data for graphs presented in Figures 3B, D, F, H, J.

| Table S2 (Figure) | Parameter | *p* - value comparison | Genotype-treatment | Mean ± SD (%) | *p* - value | n |
|---|---|---|---|---|---|---|
| Fig. 3B | calpain activity | $R^{I255d/+}$ - NT | WT - NT | 1.22 ± 0.27 | $p = 0.0184$ | 7 |
| | | | $R^{I255d/+}$ - NT | 2.74 ± 1.26 | | 10 |
| | | | $R^{I255d/+}$ - Ola 0.1 | 1.54 ± 0.37 | $p = 0.6430$ | 4 |
| | | | $R^{I255d/+}$ - Ola 1 | 1.19 ± 0.31 | $p = 0.3048$ | 3 |
| | | | $R^{I255d/+}$ - Ola 10 | 4.53 ± 0.90 | $p < 0.0001$ | 4 |
| Fig. 3D | calpain-2 | $R^{I255d/+}$ - NT | WT - NT | 1.31 ± 0.74 | $p = 0.0126$ | 3 |
| | | | $R^{I255d/+}$ - NT | 4.38 ± 0.46 | | 3 |
| | | | $R^{I255d/+}$ - Ola 0.1 | 2.96 ± 0.72 | $p = 0.0968$ | 8 |
| | | | $R^{I255d/+}$ - Ola 1 | 3.21 ± 0.75 | $p = 0.1463$ | 6 |
| | | | $R^{I255d/+}$ - Ola 10 | 4.88 ± 1.95 | $p > 0.9999$ | 6 |
| Fig. 3F | caspase-3 | $R^{I255d/+}$ - NT | WT - NT | 0.25 ± 0.09 | $p = 0.0208$ | 3 |
| | | | $R^{I255d/+}$ - NT | 2.52 ± 0.78 | | 6 |
| | | | $R^{I255d/+}$ - Ola 0.1 | 2.22 ± 0.52 | $p = 0.6210$ | 7 |
| | | | $R^{I255d/+}$ - Ola 1 | 3.96 ± 1.00 | $p = 0.0371$ | 6 |
| | | | $R^{I255d/+}$ - Ola 10 | 4.76 ± 1.52 | $p = 0.0331$ | 6 |
| Fig. 3H | rhodopsin | $R^{I255d/+}$ - NT | WT - NT | 1.99 ± 0.21 | $p < 0.0001$ | 7 |
| | | | $R^{I255d/+}$ - NT | 8.18 ± 1.23 | | 9 |
| | | | $R^{I255d/+}$ - Ola 0.1 | 8.45 ± 0.44 | $p = 0.9570$ | 6 |
| | | | $R^{I255d/+}$ - Ola 1 | 8.37 ± 1.62 | $p = 0.9827$ | 6 |
| | | | $R^{I255d/+}$ - Ola 10 | 8.17 ± 1.37 | $p > 0.9999$ | 7 |
| Fig. 3J | cone arrestin-3 | $R^{I255d/+}$ - NT | WT - NT | 16.74 ± 1.99 | $p = 0.0067$ | 6 |
| | | | $R^{I255d/+}$ - NT | 12.32 ± 1.82 | | 6 |
| | | | $R^{I255d/+}$ - Ola 0.1 | 13.52 ± 2.80 | $p = 0.7353$ | 6 |
| | | | $R^{I255d/+}$ - Ola 1 | 12.91 ± 1.71 | $p = 0.9686$ | 6 |
| | | | $R^{I255d/+}$ - Ola 10 | 11.81 ± 1.52 | $p = 0.9866$ | 5 |

**Table S3.** Effect of olaparib (Ola) lasting 4 days on the activity of PARP (PARP *in situ* assay) and calpain (calpain *in situ* assay), photoreceptor cell death (TUNEL) in outer nuclear layer (ONL) with concentration (1 µM): Quantitative data for graphs presented in Figures S1B, D, F.

| Table S3 (Figure) | Parameter | *p* - value comparison | Genotype-treatment | Mean ± SD (%) | *p* - value | n |
|---|---|---|---|---|---|---|
| Fig. S1B | PARP activity | $R^{I255d/+}$ - NT | WT - NT | 1.74 ± 0.33 | $p = 0.0002$ | 6 |
| | | | $R^{I255d/+}$ - NT | 3.74 ± 0.36 | | 4 |
| | | | $R^{I255d/+}$ - Ola 1 | 0.27 ± 0.05 | $p < 0.0001$ | 4 |
| Fig. S1D | calpain activity | $R^{I255d/+}$ - NT | WT - NT | 1.47 ± 0.26 | $p = 0.1040$ | 6 |
| | | | $R^{I255d/+}$ - NT | 2.16 ± 0.73 | | 4 |
| | | | $R^{I255d/+}$ - Ola 1 | 2.47 ± 0.50 | $p = 0.4466$ | 4 |
| Fig. S1F | TUNEL | $R^{I255d/+}$ - NT | WT - NT | 2.07 ± 0.29 | $p = 0.0227$ | 6 |
| | | | $R^{I255d/+}$ - NT | 6.79 ± 1.02 | | 4 |
| | | | $R^{I255d/+}$ - Ola 1 | 18.18 ± 2.78 | $p = 0.0003$ | 4 |

**Table S4.** Comparison of the Effect of 1 µM olaparib (Ola) lasting from post-natal (P) day 12 to P18 (4 days) and P20 (6 days) on the activity of PARP (PARP *in situ* assay) and calpain (calpain *in situ* assay), photoreceptor cell death (TUNEL) in outer nuclear layer (ONL), and ONL rows count: Quantitative data for graphs presented in Figures S1G, H, I, J.

| Table S4 (Figure) | Parameter | Genotype-treatment | Endpoint | Mean ± SD (%) | *p* - value compare to P18 | n |
|---|---|---|---|---|---|---|
| Fig. S1G | PARP activity | WT-NT | P18 | 1.74 ± 0.33 | | 6 |
| | | | P20 | 1.36 ± 0.36 | $p = 0.6081$ | 7 |
| | | $R^{I255d/+}$ - NT | P18 | 3.74 ± 0.36 | | 4 |
| | | | P20 | 3.33 ± 0.91 | $p = 0.5715$ | 12 |
| | | $R^{I255d/+}$ - Ola 1 | P18 | 0.27 ± 0.05 | | 4 |
| | | | P20 | 0.10 ± 0.02 | $p = 0.9740$ | 3 |
| Fig. S1H | calpain activity | WT-NT | P18 | 1.47 ± 0.26 | | 6 |
| | | | P20 | 1.22 ± 0.27 | $p = 0.9171$ | 7 |
| | | $R^{I255d/+}$ - NT | P18 | 2.16 ± 0.73 | | 4 |
| | | | P20 | 2.83 ± 1.16 | $p = 0.3788$ | 12 |
| | | $R^{I255d/+}$ - Ola 1 | P18 | 2.47 ± 0.50 | | 4 |
| | | | P20 | 1.19 ± 0.31 | $p = 0.1103$ | 3 |
| Fig. S1I | TUNEL | WT-NT | P18 | 2.07 ± 0.29 | | 6 |
| | | | P20 | 3.03 ± 0.77 | $p = 0.8426$ | 10 |
| | | $R^{I255d/+}$ - NT | P18 | 6.79 ± 1.02 | | 4 |
| | | | P20 | 7.28 ± 2.60 | $p = 0.9778$ | 20 |
| | | $R^{I255d/+}$ - Ola 1 | P18 | 18.18 ± 2.78 | | 4 |
| | | | P20 | 14.51 ± 4.02 | $p = 0.0518$ | 9 |
| Fig. S1J | ONL rows | WT-NT | P18 | 10.34 ± 0.31 | | 6 |
| | | | P20 | 8.46 ± 1.66 | $p = 0.0076$ | 9 |
| | | $R^{I255d/+}$ - NT | P18 | 6.02 ± 1.39 | | 4 |
| | | | P20 | 4.83 ± 1.07 | $p = 0.1810$ | 16 |
| | | $R^{I255d/+}$ - Ola 1 | P18 | 5.42 ± 0.69 | | 4 |
| | | | P20 | 5.53 ± 0.80 | $p = 0.9979$ | 9 |

**Table S5.** Effect of saruparib (Sar) on the activity of PARP (PARP *in situ* assay) and calpain (calpain *in situ* assay), photoreceptor cell death (TUNEL) in outer nuclear layer (ONL), with concentrations (0.1, 1, and 10 µM): Quantitative data for graphs presented in Figures 4B, D, F.

| Table S5 (Figure) | Parameter | *p* - value comparison | Genotype-treatment | Mean ± SD (%) | *p* - value | n |
|---|---|---|---|---|---|---|
| Fig. 4B | PARP activity | $R^{I255d/+}$ - NT | WT - NT | 1.28 ± 0.34 | $p = 0.0004$ | 5 |
| | | | $R^{I255d/+}$ - NT | 3.50 ± 1.04 | | 8 |
| | | | $R^{I255d/+}$ - Sar 0.1 | 1.92 ± 0.36 | $p = 0.0094$ | 5 |
| | | | $R^{I255d/+}$ - Sar 1 | 1.05 ± 0.67 | $p = 0.0001$ | 5 |
| | | | $R^{I255d/+}$ - Sar 10 | 0.11 ± 0.09 | $p < 0.0001$ | 4 |
| Fig. 4D | calpain activity | $R^{I255d/+}$ - NT | WT - NT | 1.10 ± 0.10 | $p = 0.5301$ | 5 |
| | | | $R^{I255d/+}$ - NT | 2.46 ± 1.25 | | 8 |
| | | | $R^{I255d/+}$ - Sar 0.1 | 3.56 ± 0.43 | $p = 0.0024$ | 5 |
| | | | $R^{I255d/+}$ - Sar 1 | 3.66 ± 1.03 | $p = 0.0015$ | 5 |
| | | | $R^{I255d/+}$ - Sar 10 | 4.45 ± 0.88 | $p = 0.0001$ | 4 |
| Fig. 4F | TUNEL | $R^{I255d/+}$ - NT | WT - NT | 2.91 ± 0.86 | $p = 0.0651$ | 6 |
| | | | $R^{I255d/+}$ - NT | 7.07 ± 2.55 | | 14 |
| | | | $R^{I255d/+}$ - Sar 0.1 | 9.67 ± 2.23 | $p = 0.2404$ | 5 |
| | | | $R^{I255d/+}$ - Sar 1 | 13.21 ± 3.61 | $p = 0.0023$ | 5 |
| | | | $R^{I255d/+}$ - Sar 10 | 22.52 ± 3.98 | $p < 0.0001$ | 4 |

**Table S6.** Effect of INO1001 (INO) on the activity of PARP (PARP *in situ* assay) and calpain (calpain *in situ* assay), photoreceptor cell death (TUNEL) in outer nuclear layer (ONL), with concentrations (0.1, 1, and 10 µM): Quantitative data for graphs presented in Figures 5B, D, F.

| Table S6 (Figure) | Parameter | *p* - value comparison | Genotype-treatment | Mean ± SD (%) | *p* - value | n |
|---|---|---|---|---|---|---|
| Fig. 5B | PARP activity | $R^{I255d/+}$ - NT | WT - NT | 1.26 ± 0.33 | $p = 0.0045$ | 5 |
| | | | $R^{I255d/+}$ - NT | 3.14 ± 0.77 | | 10 |
| | | | $R^{I255d/+}$ - INO 0.1 | 2.91 ± 1.03 | $p = 0.7807$ | 6 |
| | | | $R^{I255d/+}$ - INO 1 | 2.54 ± 1.16 | $p = 0.6953$ | 4 |
| | | | $R^{I255d/+}$ - INO 10 | 0.50 ± 0.26 | $p = 0.0011$ | 3 |
| Fig. 5D | calpain activity | $R^{I255d/+}$ - NT | WT - NT | 1.10 ± 0.11 | $p = 0.3810$ | 5 |
| | | | $R^{I255d/+}$ - NT | 2.49 ± 1.27 | | 8 |
| | | | $R^{I255d/+}$ - INO 0.1 | 2.63 ± 1.13 | $p = 0.9033$ | 6 |
| | | | $R^{I255d/+}$ - INO 1 | 3.02 ± 0.78 | $p = 0.3447$ | 4 |
| | | | $R^{I255d/+}$ - INO 10 | 2.26 ± 0.10 | $p = 0.9458$ | 3 |
| Fig. 5F | TUNEL | $R^{I255d/+}$ - NT | WT - NT | 2.91 ± 0.76 | $p = 0.0951$ | 6 |
| | | | $R^{I255d/+}$ - NT | 7.46 ± 2.53 | | 14 |
| | | | $R^{I255d/+}$ - INO 0.1 | 7.35 ± 1.50 | $p = 0.9998$ | 6 |
| | | | $R^{I255d/+}$ - INO 1 | 11.52 ± 3.45 | $p = 0.1738$ | 4 |
| | | | $R^{I255d/+}$ - INO 10 | 17.54 ± 5.69 | $p = 0.0013$ | 3 |

**Table S7.** Effect of nicotinamide (NAM) on the activity of PARP (PARP *in situ* assay) and calpain (calpain *in situ* assay), photoreceptor cell death (TUNEL) in outer nuclear layer (ONL), with concentrations (20, 200, 1000 and 2000 µM): Quantitative data for graphs presented in Figures 6B, D, F.

| Table S7 (Figure) | Parameter | *p* - value comparison | Genotype-treatment | Mean ± SD (%) | *p* - value | n |
|---|---|---|---|---|---|---|
| Fig. 6B | PARP activity | $R^{I255d/+}$ - NT | WT - NT | 1.39 ± 0.43 | $p < 0.0001$ | 5 |
| | | | $R^{I255d/+}$ - NT | 3.16 ± 0.77 | | 10 |
| | | | $R^{I255d/+}$ - NAM 20 | 2.62 ± 0.62 | $p = 0.0972$ | 6 |
| | | | $R^{I255d/+}$ - NAM 200 | 1.36 ± 0.42 | $p = 0.0001$ | 6 |
| | | | $R^{I255d/+}$ - NAM 1000 | 1.87 ± 0.50 | $p = 0.0017$ | 6 |
| | | | $R^{I255d/+}$ - NAM 2000 | 0.89 ± 0.21 | $p < 0.0001$ | 5 |
| Fig. 6D | calpain activity | $R^{I255d/+}$ - NT | WT - NT | 1.21 ± 0.33 | $p = 0.3754$ | 5 |
| | | | $R^{I255d/+}$ - NT | 2.58 ± 1.22 | | 9 |
| | | | $R^{I255d/+}$ - NAM 20 | 2.46 ± 0.71 | $p = 0.9444$ | 6 |
| | | | $R^{I255d/+}$ - NAM 200 | 3.11 ± 0.50 | $p = 0.9949$ | 6 |
| | | | $R^{I255d/+}$ - NAM 1000 | 2.67 ± 0.59 | $p = 0.9974$ | 6 |
| | | | $R^{I255d/+}$ - NAM 2000 | 2.53 ± 0.68 | $p = 0.9637$ | 5 |
| Fig. 6F | TUNEL | $R^{I255d/+}$ - NT | WT - NT | 2.58 ± 0.36 | $p = 0.0019$ | 6 |
| | | | $R^{I255d/+}$ - NT | 6.48 ± 2.36 | | 13 |
| | | | $R^{I255d/+}$ - NAM 20 | 6.82 ± 0.88 | $p = 0.8127$ | 6 |
| | | | $R^{I255d/+}$ - NAM 200 | 9.10 ± 1.30 | $p = 0.0431$ | 6 |
| | | | $R^{I255d/+}$ - NAM 1000 | 12.77 ± 2.89 | $p = 0.0002$ | 6 |
| | | | $R^{I255d/+}$ - NAM 2000 | 12.97 ± 2.05 | $p < 0.0001$ | 5 |

**Table S8.** Effect of DMSO and PARP inhibitors, including olaparib (Ola), saruparib (Sar), INO1001 (INO), and nicotinamide (NAM), on outer nuclear layer (ONL) rows counts, with different concentrations, DMSO (0.01, 0.1 and 1%), Ola, Sar and INO (0.1, 1 and 1µM), and NAM (20, 200, 1000 and 2000 µM): Quantitative data for graphs presented in Figures 7D.

| Table S8 (Figure) | Genotype | *p* - value comparison | Genotype-treatment | Mean ± SD (%) | *p* - value | n |
|---|---|---|---|---|---|---|
| Fig. 7D ONL rows | WT | NT | WT – NT | 8.46 ± 1.66 | | 9 |
| | | | WT – DMSO 0.01 | 10.18 ± 0.44 | *p* = 0.1565 | 4 |
| | | | WT – DMSO 0.1 | 9.58 ± 1.74 | *p* = 0.5510 | 3 |
| | | | WT – DMSO 1 | 9.23 ± 1.06 | *p* = 0.7289 | 4 |
| | $R^{I255d/+}$ | NT | $R^{I255d/+}$ - NT | 4.83 ± 1.07 | | 16 |
| | | | $R^{I255d/+}$ - DMSO 0.01 | 5.73 ± 0.47 | *p* = 0.3541 | 3 |
| | | | $R^{I255d/+}$ - DMSO 0.1 | 5.87 ± 0.64 | *p* = 0.2467 | 3 |
| | | | $R^{I255d/+}$ - DMSO 1 | 5.82 ± 0.17 | *p* = 0.2795 | 3 |
| | $R^{I255d/+}$ | NT | $R^{I255d/+}$ - NT | 4.92 ± 1.09 | | 14 |
| | | | $R^{I255d/+}$ - Ola 0.1 | 5.42 ± 0.82 | *p* = 0.5613 | 9 |
| | | | $R^{I255d/+}$ - Ola 1 | 5.53 ± 0.80 | *p* = 0.4001 | 9 |
| | | | $R^{I255d/+}$ - Ola 10 | 4.24 ± 1.39 | *p* = 0.4313 | 6 |
| | $R^{I255d/+}$ | NT | $R^{I255d/+}$ - NT | 4.93 ± 1.14 | | 14 |
| | | | $R^{I255d/+}$ - Sar 0.1 | 4.99 ± 0.94 | *p* = 0.9989 | 5 |
| | | | $R^{I255d/+}$ - Sar 1 | 4.48 ± 0.46 | *p* = 0.7287 | 5 |
| | | | $R^{I255d/+}$ - Sar 10 | 4.60 ± 0.38 | *p* = 0.8950 | 4 |
| | $R^{I255d/+}$ | NT | $R^{I255d/+}$ - NT | 4.93 ± 1.09 | | 14 |
| | | | $R^{I255d/+}$ - INO 0.1 | 4.22 ± 0.39 | *p* = 0.3078 | 6 |
| | | | $R^{I255d/+}$ - INO 1 | 4.37 ± 0.87 | *p* = 0.6109 | 4 |
| | | | $R^{I255d/+}$ - INO 10 | 4.44 ± 0.17 | *p* = 0.7757 | 3 |
| | $R^{I255d/+}$ | NT | $R^{I255d/+}$ - NT | 4.96 ± 1.10 | | 14 |
| | | | $R^{I255d/+}$ - NAM 20 | 5.40 ± 0.70 | *p* = 0.7414 | 6 |
| | | | $R^{I255d/+}$ - NAM 200 | 5.39 ± 0.86 | *p* = 0.7579 | 6 |
| | | | $R^{I255d/+}$ - NAM 1000 | 5.62 ± 0.55 | *p* = 0.3991 | 6 |
| | | | $R^{I255d/+}$ - NAM 2000 | 5.56 ± 0.59 | *p* = 0.5496 | 5 |
